# Characterization and Functional Annotation of Uncharacterized ABC Transporter ATP-Binding Protein Rv0986 of *Mycobacterium tuberculosis* (Strain ATCC 25618 / H37Rv)

**DOI:** 10.1101/2020.05.23.112680

**Authors:** Abu Saim Mohammad Saikat

**Affiliations:** Department of Biochemistry and Molecular Biology, Bangabandhu Sheikh Mujibur Rahman Science and Technology University, Gopalganj, Bangladesh

**Keywords:** *Mycobacterium tuberculosis*, Protein Rv0986, Homology Modeling, Ligand binding site, Tuberculosis

## Abstract

The most significant ancient infectious disease tuberculosis is causes by a human pathogen, *Mycobacterium tuberculosis* (MTB). Amazingly, tuberculosis (TB) has become one of the major causes of human death worldwide. The protein Rv0986 is associated with the ATP-binding cassette domain of the transporters involved in the export of lipoprotein and macrolide, and cell division protein, therefore, related to mycobacterial infection. But the protein Rv0986 is not yet explored. As a result, identification, characterization, and functional annotation of uncharacterized protein Rv0986 were predicted where the structure modeling was generated by using Modeller, Phyre2, and Swiss Model with the structural quality assessment by Ramachandran Plot (PROCHECK), Verify 3d, and Swiss-Model Interactive Workplace as well. Z-scores obtained from Prosa-web were also applied for overall 3D model quality. This in-silico method will uncover the significance of undiscovered uncharacterized protein Rv0986 present in MTB, and indeed it can accelerate the way to enrich our knowledge in the pathogenesis and drug-targeting opportunity against infection by MTB.

## 1 Introduction

*Mycobacterium tuberculosis* (MTB) is a Gram-positive, acid-fast, and rod-like organism which causes tuberculosis (TB). TB is one of the top lethal communicable diseases (ranking above HIV/AIDS) worldwide. It spreads from infected people with TB banish bacteria into the air; for example, by coughing. Normally, it affects the lungs (pulmonary TB) as well as other sites of the body (extrapulmonary TB). It is estimated that around 10.0 million (range: 9.0–11.1 million) people are in new TB cases and 1.2 million (range, 1.1–1.3 million) TB deaths among HIV-negative people in 2018, resulting in a high risk of TB disease development and spreading globally [1] [2] [3]. The uncharacterized ABC transporter ATP-binding protein Rv0986 of *Mycobacterium tuberculosis* (strain ATCC 25618 / H37Rv) is the ATPase catalytic subunit of an ABC transporter complex responsible for coupling the energy of ATP hydrolysis to the import of one or more from a variety of substrates including lipoproteins, macrolide, and hemin, such as lipoprotein-releasing system ATP-binding protein LolD thus enhancing mycobacterial infection [4][5][6]. However, the structure of the protein Rv0986is not reported yet. The detailed physicochemical characterization and putative structure with ligand binding active sites are not elucidated, therefore, the In Silico 3D structure prediction of Rv0986 present in MTB with characterization and functional annotation of the protein is proposed by applying *in silico* structure modeling.

## 2 Methodology

### 2.1 Retrieval of Target Amino Acid Sequence

The amino acid sequence of uncharacterized ABC transporter ATP-binding protein Rv0986 (strain ATCC 25618 / H37Rv) was obtained from UniProtKB [7] with the accession ID P9WQK1. As the 3D structure is unavailable in the Protein Data Bank (PDB), modeling of this unexplored protein was undertaken to utilize a 248 amino acid long sequence of protein Rv0986 present in MTB.

### 2.2 Physicochemical Characterization

The physicochemical properties of the retrieved sequence were determined using two web-based servers. The ProtParam tool of ExPasy [8] employed for the prediction of amino acid composition, instability and aliphatic index, extinction coefficients, and grand average of hydropathicity (GRAVY). Theoretical isoelectric point (pI) was also calculated using the Sequence Manipulation Suite (SMS) version 2 [9]. For the domain prediction of the protein Rv0986, the Conserved Domain (CD) Search Service tool of the National Center for Biotechnology Information (NCBI) [10] was used; and for the determination of motif ScanProsite Tool [11] was applied.

### 2.3 Secondary Structure Prediction

The self-optimized prediction method with alignment (SOPMA) [12] and SPIPRED [13] program was used to predict the secondary structure of Uncharacterized protein Rv0986 Disorder prediction was performed using the DISOPRED tool [14].

### 2.4 Structure Modeling and Validation

As there is no experimentally concluded tertiary structure available for ABC transporter ATP-binding protein Rv0986 (strain ATCC 25618 / H37Rv) of *Mycobacterium tuberculosis* in the Protein Data Bank (PDB), homology structure modeling was done using three programs including Modeller with HHpred tool, Phyre2, and Swiss-Model server. The designed 3D models generated from Modeller, Phyre2, and Swiss-Model were compared and the most suitable 3D model was selected for the final validation. The final modeled structure (3D) was validated using Ramachandran plot analysis (PROCHECK) and the Verify 3D (https://servicesn.mbi.ucla.edu/Verify3D/). The Swiss-Model Interactive Workplace (https://swissmodel.expasy.org/assess) was applied for the final 3D model quality assessment. Z-scores derived from the Prosa-web were applied for the overall 3D model quality validation as well.

**Table 1:** Physicochemical Parameters Computed Using ProtParam and SMS Tool.

| Physicochemical Parameters | Values |
| --- | --- |
| Number of amino acids | 248 |
| Molecular weight | 27373.11 |
| Theoretical isoelectric point (pI) | 5.63, 5.72* |
| Aliphatic index | 95.52 |
| The instability index (II) | 32.44 |
| Total number of negatively charged residues (Asp + Glu) | 30 |
| Total number of positively charged residues (Arg + Lys) | 26 |
| Grand average of hydropathicity (GRAVY) | -0.265 |

\*pI determined by SMS Version2

**Table 2:** Secondary Structure Elements by SOPMA.

| Secondary Structure Elements | Values (%) |
| --- | --- |
| Alpha helix (Hh) | 47.18 |
| 3<sub>10</sub> helix (Gg) | 0.00 |
| Pi helix (Ii) | 0.00 |
| Beta bridge (Bb) | 0.00 |
| Extended strand (Ee) | 14.92 |
| Beta turn (Tt) | 5.24 |
| Bend region (Ss) | 0.00 |
| Random coil (Cc) | 32.66 |
| Ambiguous states | 0.00 |
| Other states | 0.00 |

**Table 3:** Ramachandran Plot Analysis.

| Server | Ramachandran Plot Calculation | Value (%) |
| --- | --- | --- |
| **Modeller** | Residues in most favored regions [A,B,L] | 95.2 |
|  | Residues in additional allowed regions [a,b,l,p] | 4.8 |
| | Residues in generously allowed regions [∼a,∼b,∼l,∼p] | 0.0 |
|  | Residues in disallowed regions | 0.0 |
| **Phyre2** | Residues in most favored regions [A,B,L] | 87.9 |
|  | Residues in additional allowed regions [a,b,l,p] | 11.2 |
| | Residues in generously allowed regions [∼a,∼b,∼l,∼p] | 0.5 |
|  | Residues in disallowed regions | 0.5 |
| **Swiss Model** | Residues in most favored regions [A,B,L] | 92.9 |
|  | Residues in additional allowed regions [a,b,l,p] | 6.4 |
| | Residues in generously allowed regions [∼a,∼b,∼l,∼p] | 0.5 |
|  | Residues in disallowed regions | 0.2 |

## 3 Results and Discussion

### 3.1 Physicochemical Characterization

The amino acid sequence of the protein Rv0986 present in MTB was retrieved in FASTA format and used as a query sequence for the determination of physicochemical parameters. The instability index of the protein Rv0986 of MTB is 32.44 (<40) indicates the stable nature of the protein [15]. The protein is acidic (pI 5.63, 5.72*) with a molecular weight of 27373.11 Da. Higher aliphatic index values (95.52) of the query protein suggests as a positive factor for increased thermos-stability for a wide temperature range [16]. The hydrophilic nature of the protein and the possibility of better interaction with water [17] were indicated by the lower grand average of hydropathicity (GRAVY) indices value (−0.265) as shown in table 1.

### 3.2 Secondary Structure Prediction

The default parameters (window width: 17; similarity threshold: 8; division factor: 4) were considered by SOPMA for the secondary structure prediction. Utilizing 248 proteins (sub-database) and 33 aligned proteins, SOPMA predicted 32.66 percent of residues as random coils in comparison to alpha-helix (47.18%), extended strand (14.92%) and Beta turn (5.24%) as shown in table 2. PSIPRED showing the higher confidence of the prediction of the helix, strand, and coil (Fig.1).

**Figure 1.**
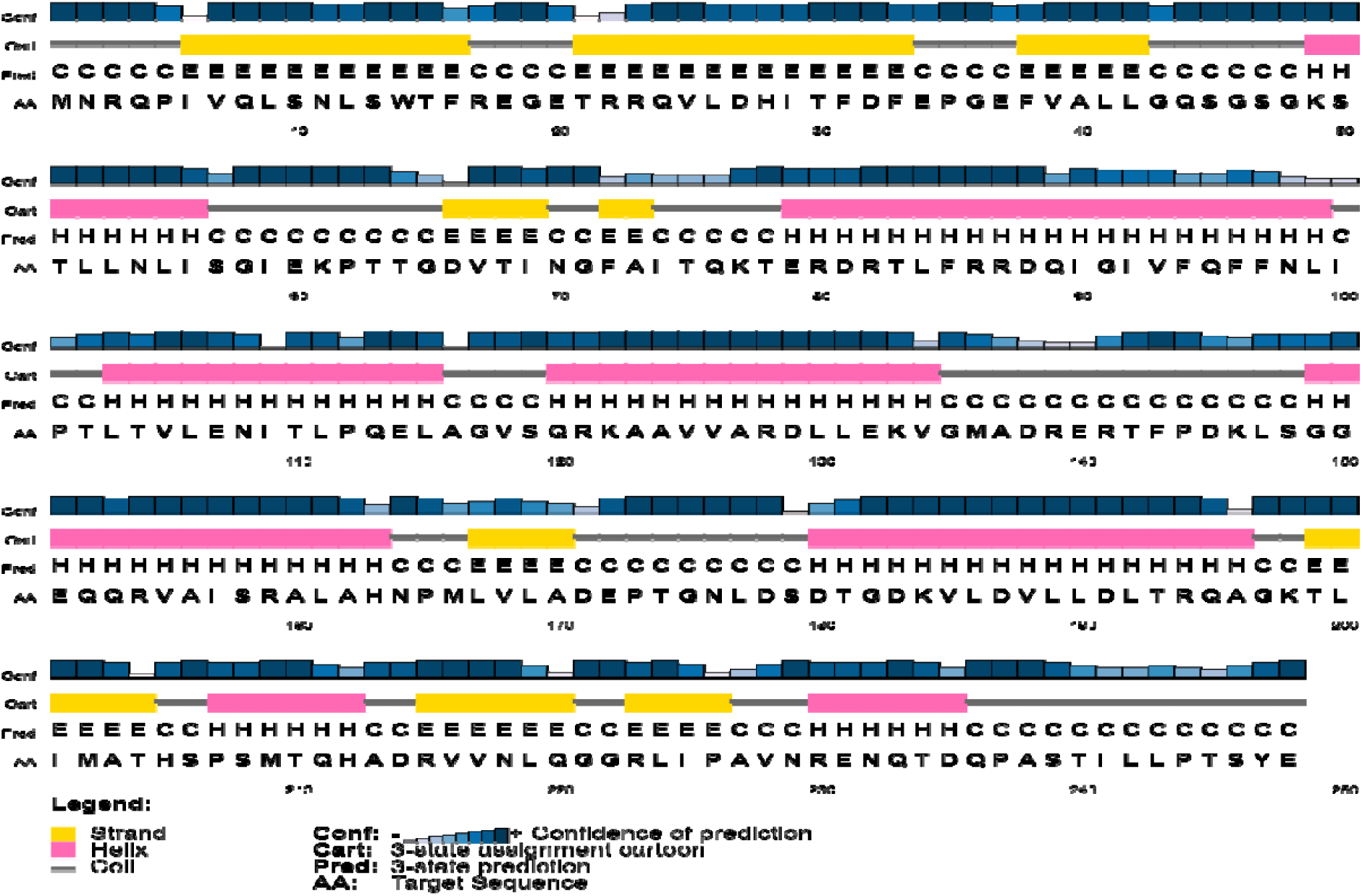
Secondary Structure Prediction by SPIPRED.

### 3.3 Protein-Protein and Protein-Polynucleotide Binding Sites

Binding sites were predicted using predict protein server, where 11 different protein binding sites were identified at positions viz.: 1-4, 17-22; 61-62; 84; 87-88; 120-122; 132-133; 139-140; 193; 212; 229-234; 246, and 4 different polynucleotide binding sites were identified at positions viz.: 45; 47; 49; and 49-50 (fig. 2).

**Figure 2.**
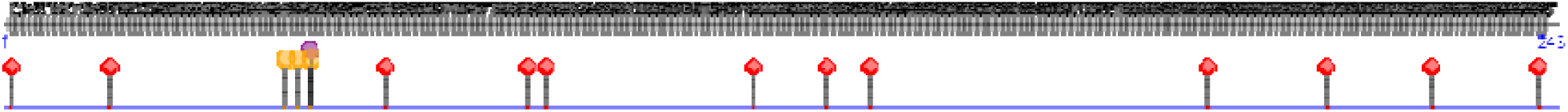
Protein-Protein and Protein-Polynucleotide Binding Sites.

### 3.4 Structure Modeling and Validation

The target sequence of the protein Rv0986 present in MTB in FASTA format was inserted to HHpred Template Selection tool [18] as input and the most effective template was selected (3TUI_D) among the number of hits of 250 with the probability rate of 100 percent, E-Value of 6.6e-32, SS of 27.5, Cols of 239 and the target length of 366 (data are not shown) and finally saved the PDB format after forwarding the submitted file to Modeller [19] (Fig. 3). The 3D structure assessment was determined by Ramachandran Map (PROCHECK) showed (Table 3) that 95.2%of the total residues were found in the core [A, B, L]; 4.8% of residues were in the additional allowed regions [a,b,l,p]; and there was no residue in the both generously allowed regions [∼a,∼b,∼l,∼p] and in the disallowed regions (fig. 5). The number of non-glycine and non-proline residues was 209 which was 100%; the end-residues (excl. Gly and Pro) were 2; the glycine residues and proline residues were 18 and 11, respectively, among the total residues(Fig. 3). The 3D model quality assessment was determined by assessment tools to Verify 3D. The 3D model passed this assessment experiment and scored 85.83% where the minimum score to pass is 80% (data are not shown). The 3D model assessment was also determined by Swiss-Model Interactive Workplace which validated this model as the MolProbity Score was 3.02 and Ramachandran favored was 97.48% with the QMEAN (Qualitative Model Energy Analysis), Cβ, All Atom, solvation, and torsion values of −1.15, −2.72, −3.06, 0.18, and −0.61, respectively (data are not shown).

**Figure 3.**
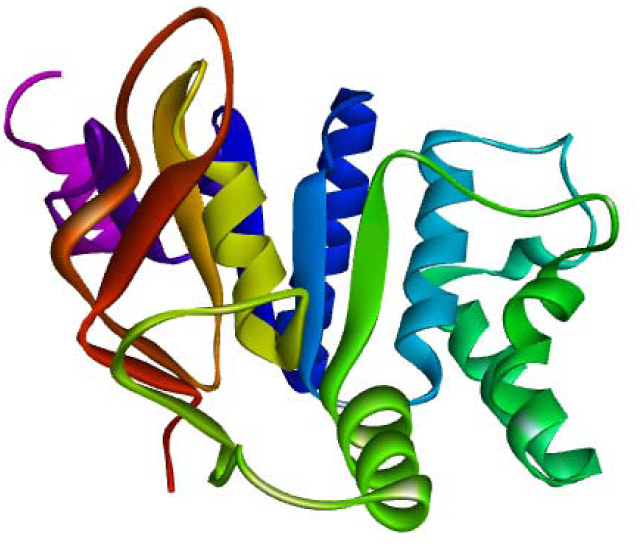
Structure of Rv0986 Predicted by Modeller.

**Figure 4(a):**
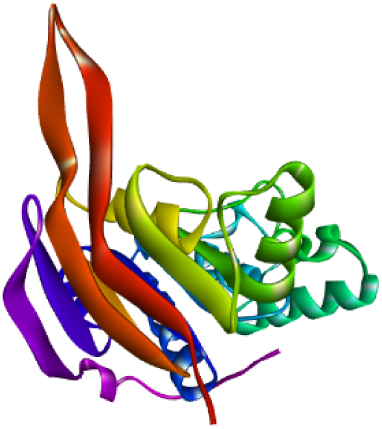
Structure of Rv0986 Predicted by Phyre2.

**Figure 4(b):**
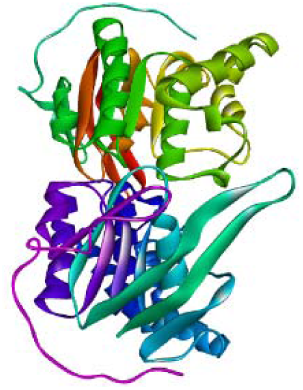
Structure of Rv0986 Predicted by Swiss Model.

**Figure 5.**
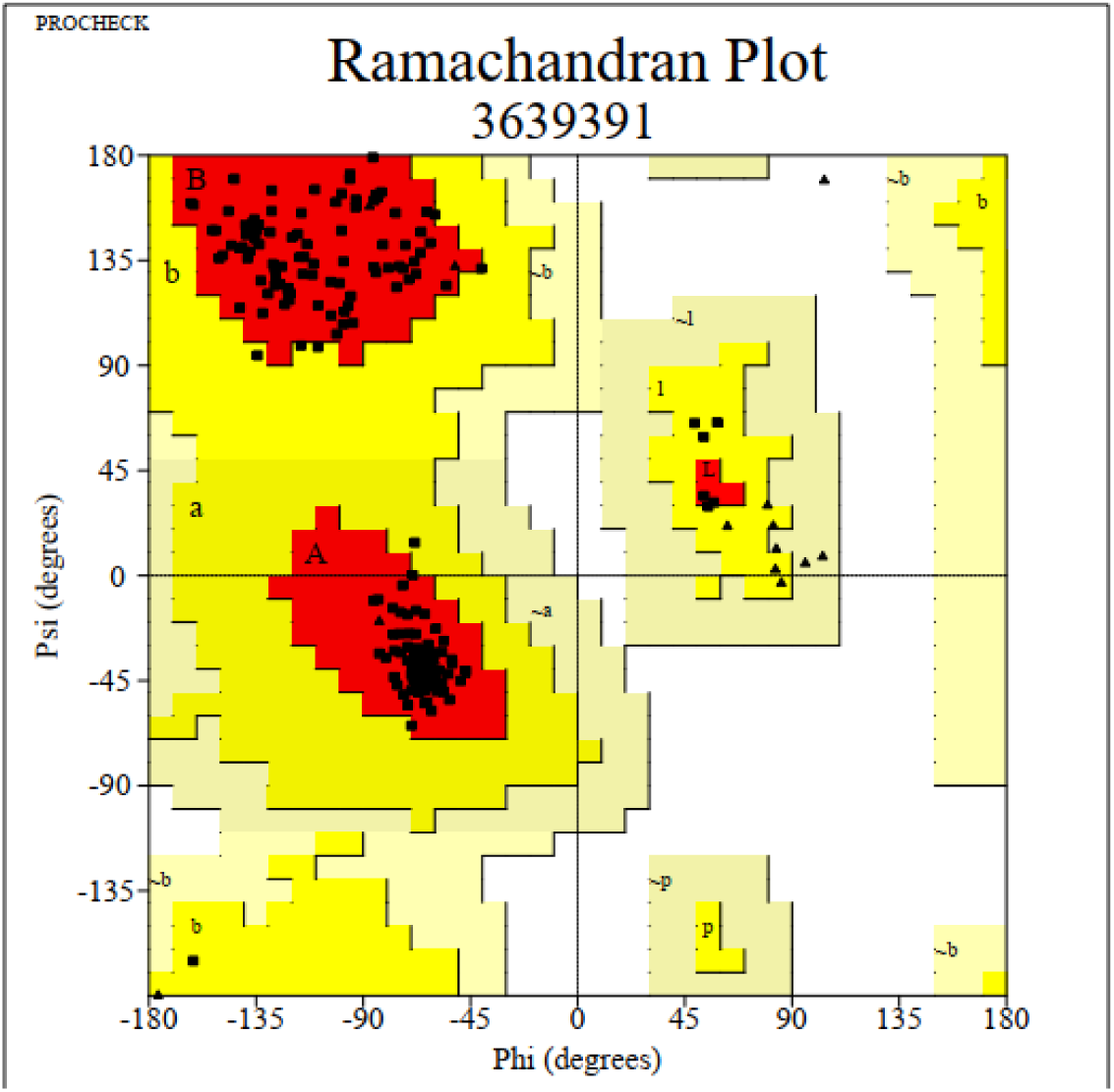
Ramachandran Plot Analysis.

Similarly, the 3D model of the protein Rv0986 was performed with Phyre2 based on the most suitable template (c5ws4A_) with the value of confidence of 100.0% and coverage of 99% (Fig. 4 -a). 245 residues (99% of the protein sequence) have been modeled with 100.0% confidence by the single highest scoring template with Phyre2. Secondary structure prediction by Phyre2 was described as the disordered of 16%, alpha-helix of 39%, and beta-strand of 25% (data are not shown). The 3D structure assessment was performed by Ramachandran Map (PROCHECK) which showed 87.9% of the residues in most favored regions; 11.2% were in additional allowed regions; 0.5% were in disallowed regions, and there was 0.5% residue in the generously allowed regions (Table 3). The 3D model quality assessment was determined by assessment tools to Verify 3D that showed the 3D model passed this assessment experiment and scored 76.42% where the minimum score to pass the quality assessment was 80% (data are not shown).

The 3D model assessment was also determined by Swiss-Model Interactive Workplace which validated this model as the MolProbity Score was 2.85 and Ramachandran favored of 91.80% with the QMEAN, Cβ, all-atom, solvation, and torsion values of −1.92, −1.81, −1.88, 0.54, and - 1.81, respectively (data are not shown) that validated the predicted 3D structure of the protein Rv0986.

The 3D model of the protein Rv0986 was also executed with Swiss-Model based on the top five suitable templates (5lj6.1.A, 5lil.1.B, 5lil.1.A, 5lil.1.B, and 2ouk.5.A) and the target sequence was selected based on the Qualitative Model Energy Analysis (QMEAN) score (−1.03), Global Model Quality Estimate (GMQE) score of 0.73, percentage of sequence identity of 39.50, and the coverage of 96%. The model was saved in PDB format when it was generated. The 3D structure assessment was determined by Ramachandran Map (PROCHECK) showed (Table 3) that 92.9% of the total residues were found in the core [A, B, L]; 92.9% of residues were in the additional allowed regions [a,b,l,p]; and there was 0.5% of residue in the generously allowed regions [∼a,∼b,∼l,∼p] and 0.2% residue was in the disallowed regions. The number of non-glycine and non-proline residues was 423 which was 100%; the end-residues (excl. Gly and Pro) were 4; the glycine residues and proline residues were 36 and 22, respectively, among the total residues (Fig. 4-b). The 3D model quality assessment was determined by assessment tools to Verify 3D. The 3D model passed this assessment experiment and scored 78.56% where the minimum score to pass is 80% (data are not shown). The 3D model assessment was also determined by Swiss-Model Interactive Workplace which validated this model as the MolProbity Score was 1.53 and Ramachandran favored was 96.88% with the QMEAN (Qualitative Model Energy Analysis), Cβ, All Atom, solvation, and torsion values of −1.03, −2.87, −1.07, 0.10, and - 0.53, respectively (data are not shown). The modeled structures of the protein Rv0986 were validated by another structure validation server, Prosa-web [20]. Standard bond angles in the modeled 3D structures were determined by the Prosa-web. Z-score was used to estimate the ‘degree of nativeness’ of the predicted structures. Z-score for the modeled 3D structure from Modeller, Phyre2, and Swiss-Model were −7.42, −7.31, and −7.37, respectively. In this paper, all three i.e. Modeller, Phyre2, and Swiss-Model servers are presenting similar values that validated the 3D structures.

## 4 Conclusion

In this study, it is concluded that the structural model of the protein Rv0986 of MTB with predicted active sites for ligand binding through *in silico* approach where the position of amino acid in the favored region by evaluating the structure. The physicochemical parameters prediction and functional annotation are useful for understanding the action of this protein’s activity. The homology-modeled protein with its predicted subcellular location in the MTB cell provides insights into the functional role of the protein Rv0986 in pathogenesis which will help to design potential therapeutic drugs against the protein present in MTB.

## Acknowledgments

I would like to thank the Department of Biochemistry and Molecular Biology of Bangabandhu Sheikh Mujibur Rahman Science and Technology University, Gopalganj, Bangladesh for providing a computational platform for completing this project.

## Declarations

### Funding

There was no funding received for this manuscript.

### Conflict of interest

The author declares no conflict of interest.

## Notes

### Competing Interest Statement

The authors have declared no competing interest.

### Summary of Updates

Updated Tables title and notes. Correspondence is added as well.

